# Moonstruck sleep: Synchronization of Human Sleep with the Moon Cycle under Natural Conditions

**DOI:** 10.1101/2020.06.01.128728

**Authors:** Leandro Casiraghi, Ignacio Spiousas, Gideon Dunster, Kaitlyn McGlothlen, Eduardo Fernández-Duque, Claudia Valeggia, Horacio O. de la Iglesia

## Abstract

As humans transitioned from hunter-gatherer to agricultural to highly urbanized post-industrial communities they progressively created environments that isolated sleep from its ancestral regulators, including the natural light-dark cycle. A prominent feature of this isolation is the availability of artificial light during the night, which delays the onset of sleep and shortens its duration. Before artificial light, moonlight was the only source of natural light sufficient to stimulate activity during the night; still, evidence for the modulation of sleep timing by lunar phases under natural conditions is controversial. Here we use data collected with wrist actimeters that measure daily sleep to show a clear synchronization of nocturnal sleep timing with the lunar cycle in participants who live in environments that range from a rural setting without access to electricity to a highly urbanized post-industrial one. The onset of sleep is delayed and sleep duration shortened as much as 1.5 hours on nights that precede the full moon night. Our data suggests that moonlight may have exerted selective pressure for nocturnal activity and sleep inhibition and that access to artificial evening light may emulate the ancestral effect of early-night moonlight.

---

Timing and duration of sleep have changed vastly throughout human evolution and history, following changes in social organization and subsistence. Human beings, with reduced vision capabilities in low-lit environments, are mostly diurnal and it is believed that nomadic groups timed their sleep onset to the time after dusk, when it became too dark to be safe hunting and gathering^1^. The establishment of industrial societies, with the availability of artificial light sources, allowed humans to accommodate their sleep and wake patterns to modern society demands by creating well-lit—or darkened—environments that isolated them considerably from natural cycles. These artificially lit environments, which can acutely inhibit sleep, entrain the central body clock in the brain that controls the timing of sleep leading to a delayed onset of sleep and a shorter nocturnal sleep bout^1–3^.

While the sun is the most important source of light and synchronizer of circadian rhythms for almost all species, moonlight also has the ability to modulate nocturnal activity in organisms ranging from invertebrate larvae to primates^4^. However, whether the moon cycle, and more specifically moonlight associated with the 29.53-day long synodic month, can modulate human nocturnal activity and sleep remains a matter of controversy. Whereas some authors have argued against strong effects of moon phase on human behaviour and biological rhythms^5,6^, recent studies have reported that human sleep and cortical activity under strictly controlled laboratory conditions, which are absent from external temporal cues, are synchronized with lunar phases^7,8^.

A full moon night is so bright to the human eye that it is indeed difficult to imagine that, in the absence of other sources of light, moonlight could not have had a role in modulating human nocturnal activity and sleep. To examine this hypothesis, we conducted a study with the Western Toba/Qom communities of the Argentinian province of Formosa. Once exclusively hunter-gatherers, these geographically spread indigenous communities share a recent historical past and live under very different levels of urbanization^9^. Some are settled in rural, isolated environments without electricity while others have access to electricity, consolidated roads and street lighting. We tested the prediction that, in communities without access to electricity, moonlit nights would be associated with increased nocturnal activity and decreased sleep.

We worked with three Toba/Qom communities (see detailed information in the Methods and Extended Data Table 1): one in an urban setting with full access to electricity (Ur), and two rural communities, one with access to limited electric light (Ru-LL) and another with no access to electric light at all (Ru-NL). Consistent with previous studies^2^, a linear mixed-effects model (LMEM) including community, sex, and age as fixed effects and subject identity as random factor revealed an association between sleep duration and onset of sleep with increased access to electric light (Extended Data Figure 1 and Extended Data Table 2). Mean sleep duration—defined as the length of the nocturnal bout of sleep—in the Urban community was 34 [95%CI: 20-48] min (Cohen’s D = 1.103) and 22 [8-36] min (D = 0.704) shorter than in the Ru-NL and Ru-LL communities, respectively. The mean onset of sleep was delayed in the Ur relatively to the Ru-NL community by 22 [4-40] min (D = 0.525). Both the duration of sleep and the time of sleep onset showed a clear modulation throughout the moon cycle that was evident in the whole population, as well as in the individual communities (Figure 1.A-D). Although intuitively one would expect sleep to be more disrupted during full moon nights, peaks and troughs for both variables did not quite match the synodic full and new moon phases but seemed to precede them by 3-5 days.

**Figure 1.**
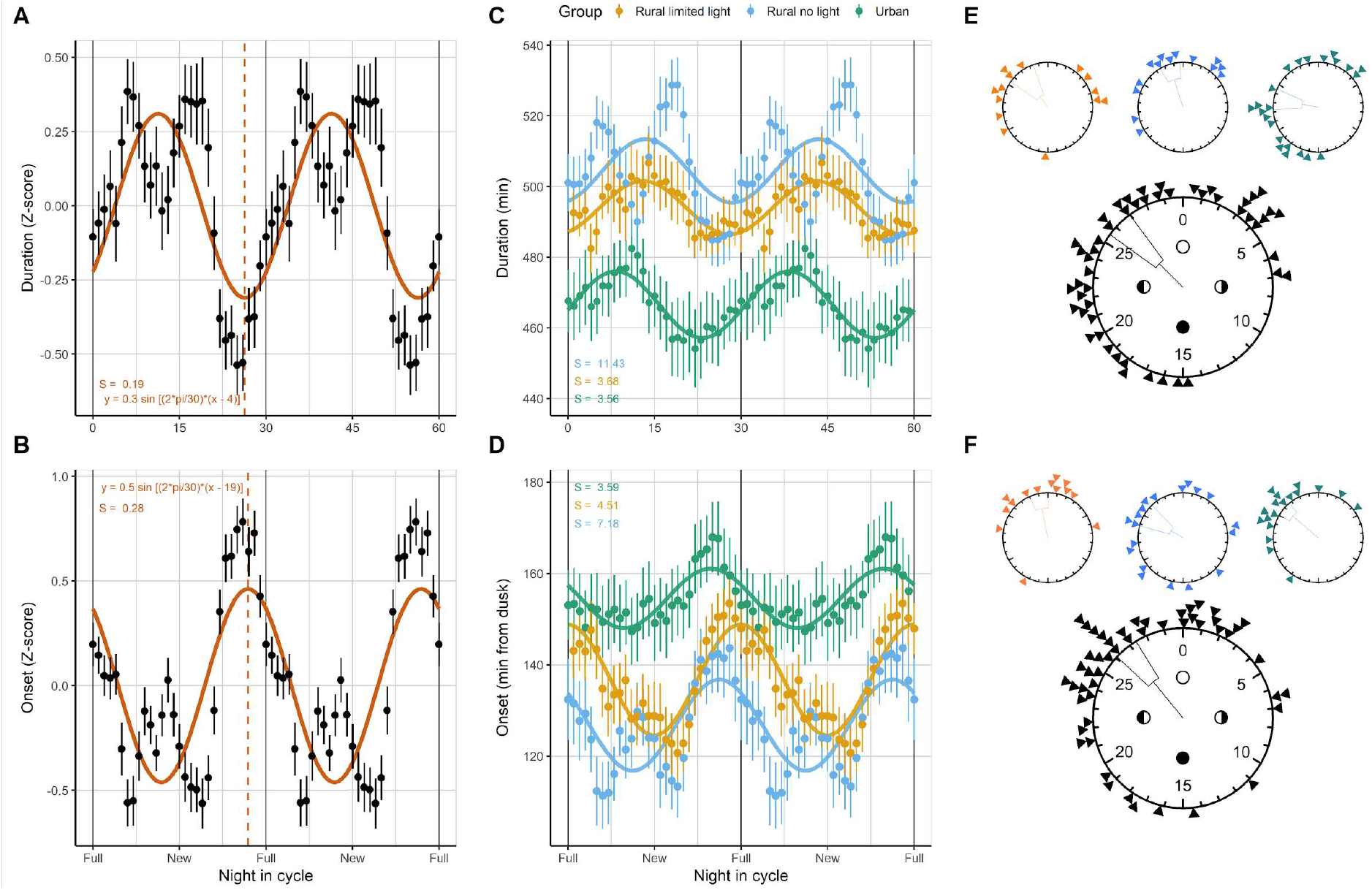
Association of sleep duration and onset with the moon cycle. **A, B)** Double-plots of the average duration and onset of sleep (measured from the time of astronomical dusk) in the Toba/Qom population across the moon cycle expressed as average z-scores (±SEM; N = 69 participants). Solid lines represent the best-fit to sinusoidal curves with a 30-day period from a non-linear least-squares fit. The best-fit sinusoid is plotted for each case, and the vertical dashed lines indicate the trough of sleep duration (i.e. the day of shorter sleep events) and the acrophase of sleep onset (i.e. the latest sleeping times). **C, D)** Double-plots of the average values of duration and onset of sleep by community (±SEM; participants: Ru-LL, 20; Ru-NL, 23; Ur, 26). **E, F)** Circular distribution of the troughs of sleep duration and acrophases of sleep onset from subjects in the three better quartiles of individual fits. The direction of the vectors indicates the mean calculated phase in days from the full moon, and the brackets the standard error of the mean. Triangles represent individual phases. Mean phase [fiducial limits] and Rayleigh z-test p-values: Duration, 26.2 [25-27.5], p = 0.0006; Onset, 26.7, [25.6-27.8], p = 3×10^−7^. Communities are split in the top row of the panels (no relevant differences between groups; see Extended Data).

The examination of individual sleep data confirms the population pattern. The modulation of sleep duration and sleep onset was evident at the individual level for participants in the three groups, in most cases displaying clear sinusoid patterns across the moon cycle (Figure 2, and Extended Data Figures 2 and 3). We selected subgroups of the subjects displaying the better fits for each variable (i.e. the three lower quartiles of the values of the standard error of the regression; 51 participants) and analysed their fit parameters. Rayleigh z-tests of the phases of the subjects in this selection indicate a consistent clustering of the troughs of sleep duration and the peaks of sleep onset on the days before the full moon (mean phase in days before the full moon [fiducial limits]: Duration, 2.8 [5.0-2.5], p = 0.0006; Onset: 3.3 [4.4-2.2], p = 3×10^−7^; Figure 1.E and F). Interestingly, some participants appeared to be in very different phases compared to the population mean—in some cases close to 180° out of phase—suggesting individual differences in the potential synchronization with the lunar cycle. Changes in sleep duration with the lunar cycle across communities, calculated from individual sinusoid fits, ranged from around 20 minutes to more than 1.5 hours and did not differ considerably between groups (mean duration change in minutes [95%CI]: Ru-NL, 46 [36-56]; Ru-LL, 52 [41-63]; Ur, 58 [50-67]). Changes in the onset of sleep varied from a handful of minutes to over an hour (Ru-NL, 29 [17-41]; Ru-LL, 32 [20-43]; Ur, 32 [24-40]).

**Figure 2.**
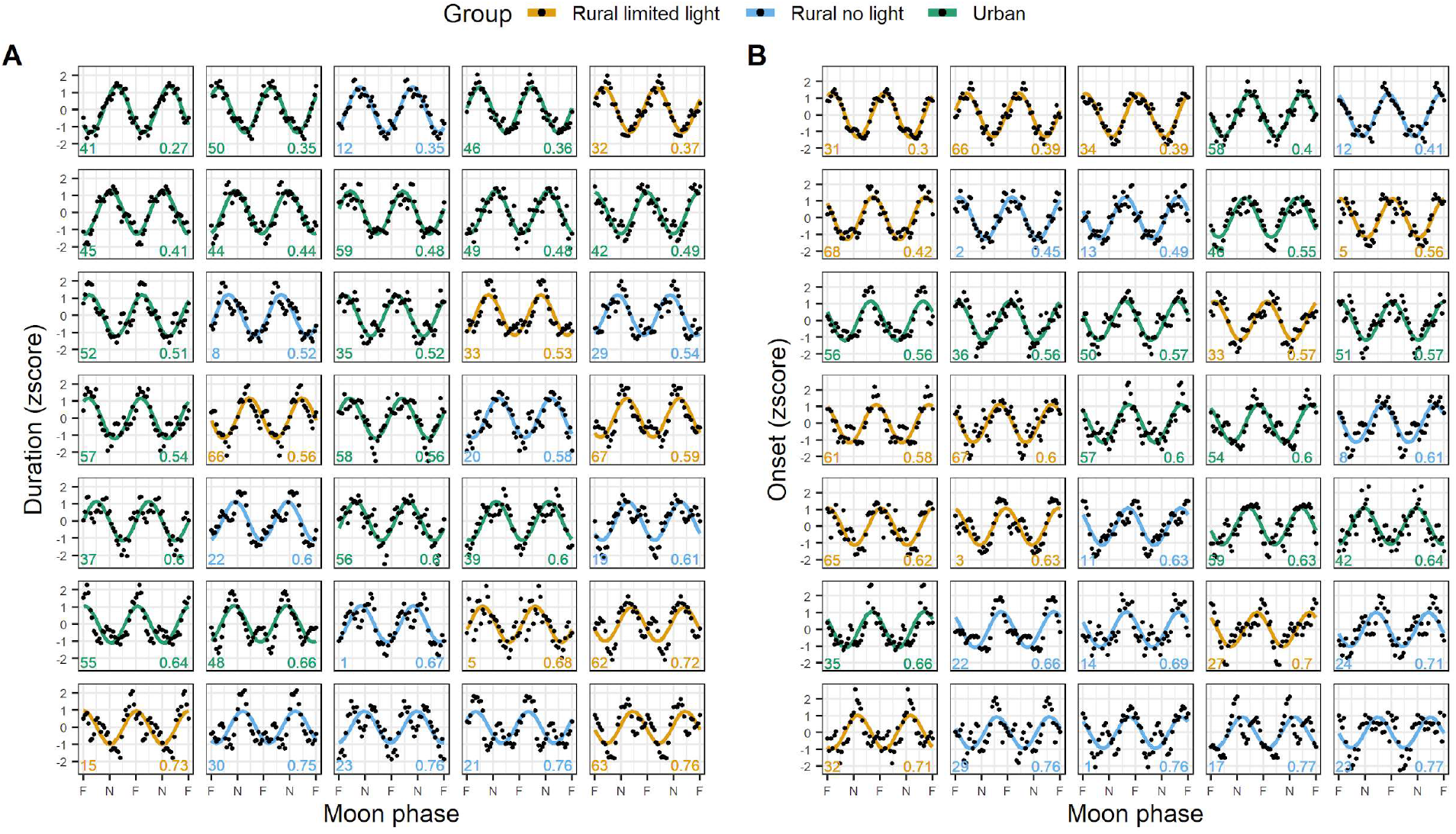
Double-plots of **A)** sleep duration and **B)** onset of sleep, expressed as z-scores, of the 35 best-fitting participants in the study for each variable. Dots indicate the data on a given night in the cycle, and coloured lines represent the best fit of a sine wave with a 30-day period through a non-linear least-squares approach. Individual data series were filtered through moving-average with a window of seven days before summarizing the data. All these data correspond to participants with records for at least 80% of the moon cycle. The number on the bottom left of each plot identifies the participant and the one on the bottom right the standard error of the regression (S). The complete set of participants are presented in Extended Data Figures 2 and 3.

Times of sleep offset showed a subtle variation across the moon cycle as well (Extended Data Figure 4.A), with later waking times preceding the full moon. Subjects in the rural community without any access to light woke up about an hour after the astronomical dawn, while the communities with access to electric light did so a little less than two hours after dawn. (Extended Data Figure 4.B).

The moon is responsible for several environmental cycles but its lighting power during the night is arguably the most relevant cycle to humans in natural conditions. Humans typically start their daily sleep bout some hours after dusk but rarely wake up before dawn, a pattern we also documented earlier among the Toba-Qom^2^. In this context, it is primarily moonlight available during the first hours of the night that is more likely to drive changes in the onset of sleep. In contrast, moonlight late in the night, when most individuals are typically asleep, will have little influence on sleep onset or duration. The hours of available moonlight change predictably through the moon cycle according to the time of moonrise (Extended Data Figure 4); under a mostly symmetrical photoperiod like that of the spring season moonlight becomes less available during the early night on the nights that follow the full moon night. We hypothesized that the adaptive value of the synchronization between sleep and the moon cycles is to stimulate wakefulness when moonlight is available during the early night, which are not necessarily the nights of brightest moonlight.

To test the predictions of this hypothesis, we determined the availability of moonlight during the first six hours of each night recorded in the study, and classified them into three categories of “*moonlight phases*” (see Methods below): Full Moonlight (F-ML), No Moonlight (No-ML), and Waning/waxing Moonlight (W-ML). We filtered the subjects with at least four data points for each moonlight phase and analysed the changes in six sleep variables. Three sleep timing variables: duration, onset and offset of sleep; and three variables that reflect sleep quality: total sleep time (the actual sleeping time within the nocturnal sleep bout), waking time after sleep onset (WASO), and fragmentation of sleep (number of waking events divided by total sleep time). Figure 3.A shows the variation of these variables according to the moonlight phase for the different communities. Within-subject averaged data comparing the F-ML and No-ML phases for different sleep parameters are shown in Figure 3.B and the summarized data is presented in Extended Data Table 3. Linear Mixed-Effects Models considering community, moonlight phase and the interaction between these, age and sex as fixed effects, and subject identity as a random effect were fit to analyse the associations between changes in sleep variables and moonlight phase (see Extended Data material for a description of the model and R syntax). According to the model, the duration of sleep for the three communities was 25 [95%CI: 13-37] minutes longer during No-ML than F-ML nights (Cohen’s D = 0.18). Accordingly, participants had 22 [12-33] more minutes of total sleep time (D = 0.18) and fell asleep 22 [13-30] minutes earlier (D = 0.21) on No-ML nights. Time of sleep offset (D = −0.029), WASO (D = 0.078), and sleep fragmentation (D = −0.040) did not show relevant differences between phases. The slopes for the interaction term between moonlight phase and community in the fitted models of all variables included the zero in the span of their confidence intervals, indicating that the effect of moon phase was similar across the three communities. The full description of the estimates and effect sizes for all factors included in the model is presented in Extended Data Table 4.

**Figure 3.**
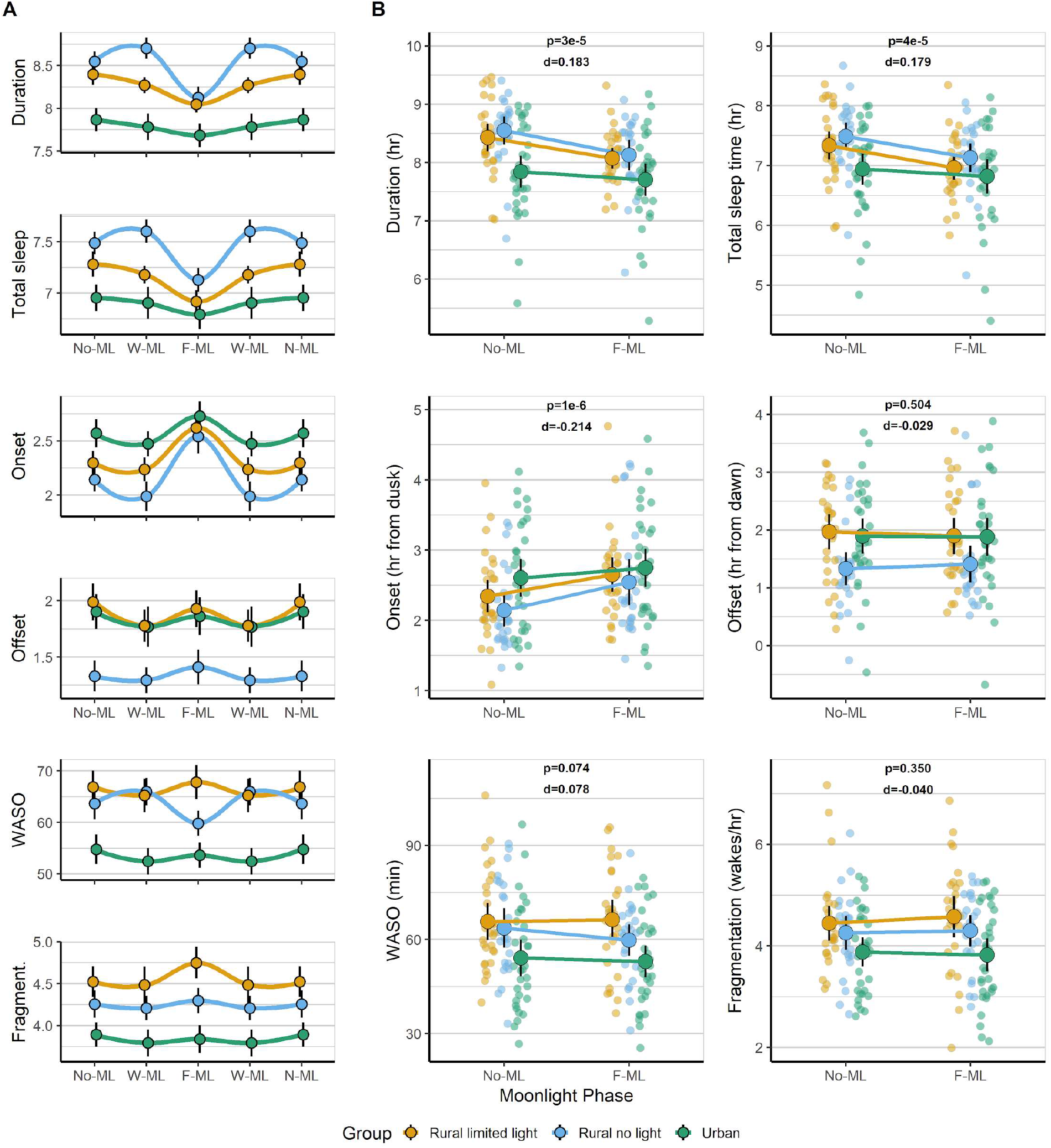
Association between the moonlight phases and sleep variables. **A)** duration of sleep, total sleep time, onset and offset of sleep, waking time after sleep onset (WASO), and fragmentation of sleep (number of wakes/total sleep time) through the “moonlight cycle” for the three communities. Data is shown as a partial double-plots displaying group means (±SEM) in the three defined moonlight phases. Connecting lines were added for illustration aid using a Local Polynomial Regression fit. **B)** Analysis of changes in the aforementioned sleep variables according to the No-ML (No Moonlight) and F-ML (Full Moonlight) phases. Smaller dots represent individual participants’ means, while the bigger dots represent the means for each community and are connected by colour lines. Vertical black error bars represent the 95% CI of the mean. The p- and Cohen’s d values for the *moonlight phase* factor in LMEMs are shown for each variable. Only participants who presented at least four data points for each moonlight phase were considered for the analysis and plots. Participants: Ru-NL, 25; Ru-LL, 25; Ur, 32.

Earlier sleep offsets did not coincide with nights in which moonlight was available during the late night, before dawn (Extended Data Figure 5). This suggests that the availability of moonlight in the late night is unlikely to drive wakefulness. To specifically explore this possibility, we compared sleep offsets between nights with and without moonlight during the six hours before dawn fitting an LMEM with the same design as in the previous moonlight phase analysis. Offset times in these two moonlight phases did not differ (see Extended Data Tables 5 and 6). Together, these results suggest that the synchronization of sleep offsets with the moon cycle is likely a by-product of the later sleep onsets during specific phases of the cycle.

Beyond moonlight phases within the ~29.5-day synodic month, the gravitational pull of the moon on the earth surface is maximal twice as frequently (every 14.75 days), during full or new moons. Recent evidence indicates that bipolar patients have mood^10,11^ and sleep (one reported case^12^) synchronized with these ~15-day semilunar phases. If a similar synchronization pattern exists with the sleep parameters of our Toba/Qom participants, we would expect that some individuals have bimodal peaks (~every 15 days) or the opposite phase relationship in their sleep parameters. Indeed, the distribution of sleep duration and onset in the Toba/Qom communities (Figure 1) as well as the inspection of the individual recordings (Extended data Figure 2) suggest the two parameters show respectively a second trough and peak within the lunar month. To further examine this possibility, we fitted sleep duration and sleep onset data to a combination of a sine wave with a 30-day period with a 15-day period harmonic wave, which led to better fitting scores (Extended Data Figures 6-8) than those of single 30-day sine waves (Figures 1–2, and Extended Data 2-3).

All three communities of Toba-Qom, including those in the urbanized setting, showed a strong association of sleep timing with the moon cycle. To explore whether a similar modulation of sleep across the moon cycle occurs in people living in large modern urban environments, we analysed sleep recordings previously obtained from over 500 University of Washington undergraduate students for a separate unpublished study. Surprisingly, the changes in sleep duration throughout the moon cycle resembled those of the Toba-Qom people, with sleep events becoming shorter in the week prior to the full moon (Figure 4). Even with the limitations inherent to observational studies, the data suggests sleep changes across the moon cycle may be still present in completely urbanized environments, where individuals may have little awareness of the synodic moon phase. In fact, light pollution measurements in the highly urbanized areas of Seattle where students typically live, reveal values that are above our full moon light measurements in the Toba/Qom rural environments. These results are also in line with two retrospective analyses of electroencephalographic recordings of sleep obtained in the controlled conditions of sleep laboratories, which found that both polysomnographic sleep and cortical activity were synchronized with lunar phases^9,10^, and point to the importance of longitudinal studies to determine the extent to which the modulation of sleep by the moon cycle prevails under modern life conditions.

**Figure 4.**
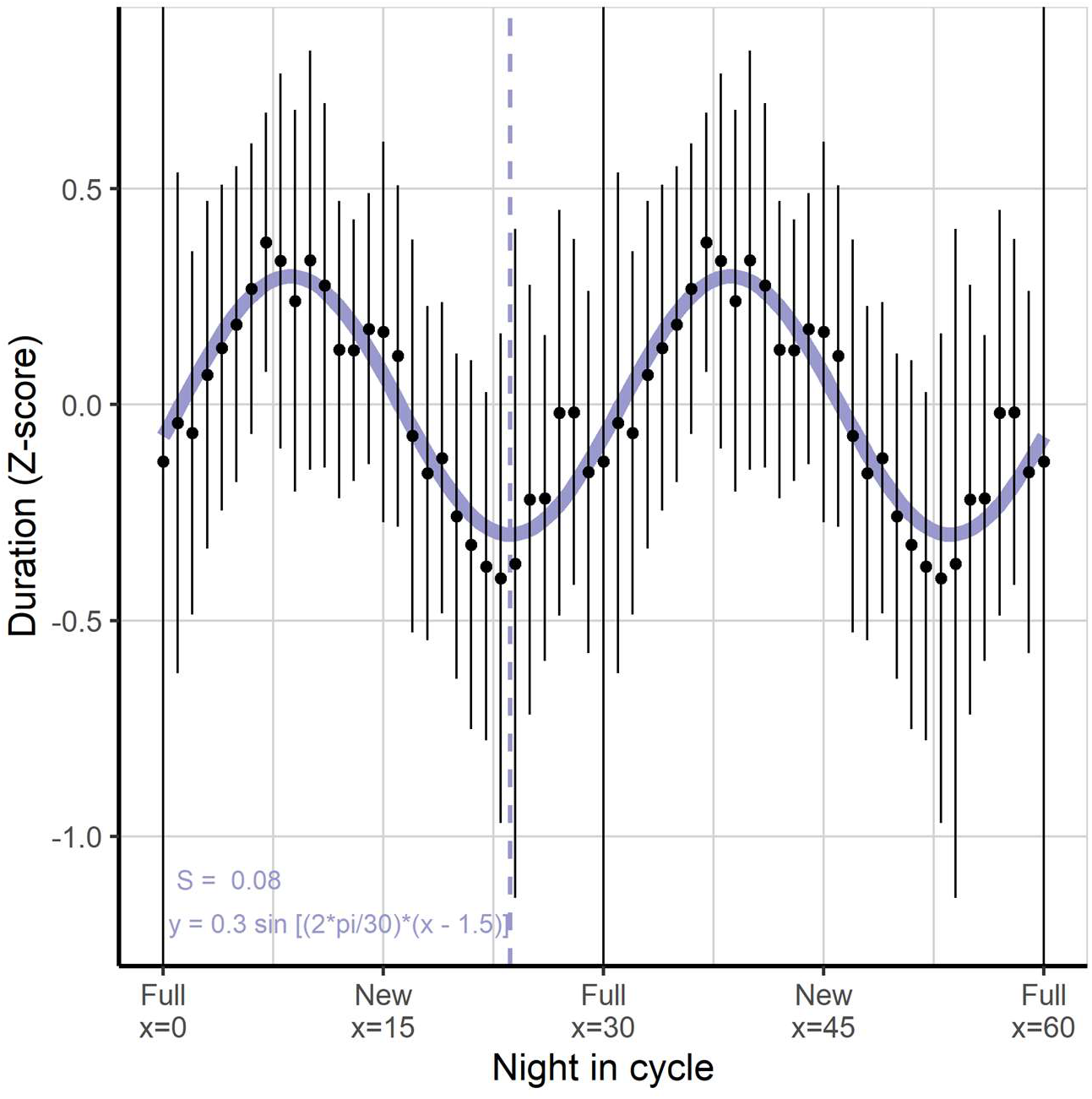
Association of sleep duration with the moon cycle in a highly urban setting. Double-plot of sleep duration (as z-scores) on weeknights recorded on 521 college students in different quarters from 2015 to 2018. The difference between individual data points and the mean values of sleep duration for weeknights in each season was calculated for over ~4,900 sleep events. The solid line represents the best sine-wave curve fit to the summarized data from a Non-Linear Least Squares fit, while the dashed line indicates the trough of this fit. The equation and standard error of the regression (S) are indicated for the fit. The complete data summary was filtered through a moving-average with a window of seven days. Error bars represent standard errors of the mean calculated individually for each night in the cycle before filtering. Number of participants/number of sleep events per quarter: Spring, 185/1835; Summer, 99/918; Fall, 143/1300; Winter, 94/840.

## Discussion

Our results show that nocturnal activity and sleep are synchronized with the moon cycle under natural conditions. Toba/Qom participants slept less and stayed up later on the days previous to full moon nights, when moonlight is available during the early night. This pattern could represent a response to the availability of moonlight during the first half of the night for communities with limited or no access to electric light. However, we were able to corroborate this modulation both in a third Toba/Qom community living with full access to electricity, as well as in a sample of college students living in a modern city. Together, these results strongly suggest that human sleep is synchronized with lunar phases regardless of ethnic and sociocultural background, and of the level of urbanization.

Increased level of access to artificial light in the Toba/Qom communities correlated with later sleep onsets and shorter duration of sleep. These findings are consistent with previous work from several laboratories including ours ^2,13–15^, as well as with the hypothesis that electric light allowed humans to extend their evening activity and push sleep times later into the night, therefore reducing the total amount of night sleep^16^. Interestingly, the availability of electric light during the evening mimics the sleep-inhibiting effects of moonlight. This is particularly evident during nights with high moonlight availability during the early hours of the night, in which the timing of sleep is most similar across the Toba-Qom communities with and without access to electric light (Fig.3.A). This finding may indicate that the effect that electric light has—delaying sleep onset and shortening sleep—could be emulating an ancestral effect of moonlit evenings, although the light intensities we are typically exposed to in our artificially lit environments are much higher.

While some studies have found minimal, or no association, between the moon cycle and sleep parameters^6,17,18^, they compared sleep during nights around the full moon to sleep during the nights of both the waxing/waning phases and the new moon. Our data clearly show these phases do not correspond to the peak and trough of sleep duration and onset. In fact, inspection of the college students’ data according to full and new moon phases, as these previous studies did, shows no clear association with sleep duration (Figure 4). In contrast to these studies and in line with our findings, two other studies found an association between sleep parameters and moon phases. Röösli et al.^19^ found an effect of moon phase on subjective sleep duration as measured by sleep diaries with shorter sleep durations around new moon nights. Similarly, in a retrospective analysis of polysomnographic sleep in a sleep lab, Cajochen et al.^7^ found an effect of moon phases on sleep duration similar to what we found at the population level, as well as on the percent of time spent on specific sleep stages.

What could be the potential adaptive value of increased activity during moonlit nights? Our interviews with Toba-Qom individuals clearly indicate moonlit nights are particularly rich in social activities. Toba-Qom elders report that at times when all sources of food came from hunting and gathering, moonlit nights had particularly high hunting and fishing activity. Furthermore, mythological stories associate the moon with the female reproductive cycle and sexual relations. The moon in the Toba culture is represented as a man who has sexual relations with women, it induces the first menstruation and regulates the timing of the following menstruations^20^. Interestingly, stories told by elder Toba/Qom point to moonlit nights as nights of higher sexual activity. These latter stories point to the possibility that ancestrally moonlight-associated encounters could have synchronized reproductive activity with women’s fertility—see the article by Helfrich-Förster et al. in this issue. Although the true adaptive value of human activity during moonlit nights remains to be determined, our data clearly indicate that humans—regardless of how urban of an environment they live in—are more active and sleep less when moonlight is available during the early hours of the night, and suggest that the effect of electric light on modern humans may have tapped into an ancestral regulatory role of moonlight on sleep.

Although our results point to moonlight availability during the early night as a likely determinant of later sleep onset and shorter sleep duration, the presence of similar lunar rhythms in sleep parameters in Seattle college students who may not be aware of the availability of moonlight, together with the presence of semilunar (~15 day-long) components on the sleep parameters of the Toba/Qom communities suggest other physical phenomena associated with the moon cycle could influence sleep. It is thus conceivable that although the ultimate cause—which confers adaptive value—for nocturnal activity in synchrony with the moon cycle is to display activity during moonlit nights, the proximal cause—which induces changes in sleep parameters—for sleep modulation by the moon cycle is the gravitational pull by the moon, which is a more reliable indicator of moon phase than its associated nocturnal illuminance.

A limitation of our observational study is that we cannot establish causality. It would be difficult to manipulate human exposure to the light the moon reflects and virtually impossible to manipulate the exposure to the gravitational pull it exerts on earth. Nevertheless, it is hard to conceive that the conserved synchronization between sleep and the moon cycle we report occurred by chance.

## Methods

### Participants and study groups

All described study procedures were approved by the Internal Review Board of University of Washington’s Human Subjects Division and were in agreement with the Declaration of Helsinki. Oral consent was obtained from every participant from the Toba-Qom communities after a verbal explanation of all procedures in Spanish. All participants were bilingual (Toba-Qom/Spanish). Parental oral consent was obtained for participants under 18 years old, who also gave their assent to participate.

Toba-Qom participants (N = 98, females 56%, mean age [range] = 24.1 ([12-75]) lived in one of three Toba/Qom communities in the Formosa province, north of Argentina. Each community had different levels of access to electric light:

1. A community located in the outskirts of Ingeniero Juárez (23°47’ S, 61°48’ W), a town with 19,000 inhabitants. All the participants in this community had 24-hour access to electric light at home and outside through streetlights, as well as to other urban features (consolidated roads, public leisure spaces, commercial premises). This community is referred to as the *Urban* group (Ur, n = 40).
2. A community located in Vaca Perdida (23°29’S, 61°38’ W), a small rural settlement of approximately 300 people 50 km north of Ingeniero Juárez. These participants had 24-hour access to electricity at their homes and light fixtures were limited at most to one light bulb per room. In contrast to Ingeniero Juárez, this community had no electric light poles in outside areas during dark hours. This community is referred to as the *Rural, limited light* group (Ru-LL, n = 33).
3. A group of participants living in sparsely distributed houses in a region known as Isla García, approximately 3 km away from Vaca Perdida. These participants lived in small extended-family groups, without any organized settlement features and no access to electric light. While children, and some adults, may have been exposed to artificial light at school, or other settings away from their home location, this could have only occurred during natural daylight times. This community is referred to as the *Rural, no light* group (Ru-NL, n = 25).

Age and sex characteristics of the three study communities are presented in Extended Data Table 1. The three study communities were similar with regard to sex (*X^2^* (2) = 0.686, p = 0.710) and age (ANOVA F (2, 96) = 0.774, p = 0.464). As expected, women displayed onset times that were 15 [95%CI: 0-30] minutes earlier than men (Cohen’s D = 0.432)^21^. Age was associated with shorter sleep duration (D = −1.411) and later sleep onset times (D = 0.429), which would be expected for the age range of our participants (Extended Data Table 2)^22,23^.

The three communities share the same ethnic and historical past^9^. In fact, the community at Ingeniero Juárez originated from a group of Toba people who migrated from the northern region in the 1990s. Housing, daily chores, and social behaviour are very similar between the communities, and the vast majority of adults are typically unemployed and rely on government subsidies.

Data were recorded during field campaigns in three consecutive years: September-October 2016 and 2017, and October-November 2018. The three campaigns were carried out during the spring (September-November in the southern hemisphere) in order to keep environmental factors (including weather, sunrise and sunset times, sunlight intensity) as stable as possible.

We also analysed data from 521 college students (mean age [range] = 21.4 [17-38], 63% females) at the University of Washington (Seattle, WA) that was recorded across different quarters between 2015 and 2018 as part of a separate study. This sample represents the portion of a total 558 students recorded who presented sleep records for at least five weeknights.

### Locomotor activity and sleep recording

Participants were equipped with Actiwatch Spectrum Plus wrist locomotor-activity loggers (Respironics, PA) for one to two months in the case of the Toba-Qom participants, and from one to three weeks for the college students. The data acquisition interval was set to one minute. Recorded data were downloaded and exported using the Philips Respironics Actiware software V.6.0.9. Participants also completed a sleep log throughout their participation, indicating times and locations of sleep events (including naps), and whether they left their home community on any given day. These logs were used for the validation of the data, and in order to discard data from time-ranges when the subjects were under different conditions to those in their main study group. The Actiware software default algorithm was used to determine the following sleep features: duration of the sleep event, total sleep time, time of sleep onset and offset, number of waking events, and waking time after sleep onset (WASO). A variable measuring of fragmentation of sleep was calculated by dividing the number of waking events during the sleep period by the total sleep time.

### Data treatment and analysis

Preparation and treatment of raw recordings, as well as the statistical analysis and plotting of data, was performed in *R* (R Core Team 2018) unless otherwise indicated.

#### Astronomical data

Sun and moon data for the Ingeniero Juárez region through the dates of recording was obtained from NASA’s JPL Horizons Web-Interface (https://ssd.jpl.nasa.gov/horizons.cgi), with a one-minute precision. The synodic moon cycle which determines the well-known “full”, “new”, and “waning” or “waxing” moon phases has an approximate average duration of 29.5 days. All nights in our study were numbered according to their position in the synodic cycle: full moon nights were considered as “night zero” with all previous and subsequent nights ordered from 1 to 29 or 30, depending on the day of the next full moon. In short, a night in position one in the cycle represents the night immediately after a full moon.

We use a second classification of phases through the moon cycle according to the availability of moonlight during the evening hours, which we refer to as “moonlight phases”. We calculated the time in the first six hours after astronomical dusk—the time that marks the disappearance of natural daylight—during which the moon was above the horizon for every night of recording, rounded this time to the nearest hour, and classified nights into three moonlight phases: 1) nights in which there was no moonlight through the first six hours of the evening (*No MoonLight*, No-ML); 2) nights during which there was moonlight available throughout the whole six hours (*Full MoonLight*, F-ML); and 3) all other nights in which the moon either rose after astronomical dusk or set before six hours after astronomical dusk (*Waning/waxing MoonLight*, W-ML). The distribution of these moonlight phases show correspondence with the standard moon phases, with the F-ML nights preceding the full moon by one week (Extended Data Figure 5.A).

#### Sleep measurements

Actograms from individual recordings were visually inspected to check for data integrity before any analysis. Low-quality criteria included: fragmented recording or very frequent missing-data periods, sustained saturated Actiwatch counts, or total lack of activity counts during extended periods of time. Seven Toba-Qom participants who displayed very low-quality recordings, due to non-compliance wearing the watch or to Actiwatch malfunction, were removed from the sample at this point.

In order to detect and discard very long and very short sleep events (artefacts from Actiwatch recording or the software algorithm), as well as irregular sleep events (long daytime siestas, naps, *all-nighters*), we analysed our sleep databases for outliers in two steps: first, looking at values of duration, and then at times of sleep onset according to the time of dusk. We performed the analyses through the median absolute deviation (MAD) method setting a threshold of 3 MADs, using the *Routliers* package for R (Leys et al. 2018).

To explore the patterns of sleep duration and timing through the moon cycle in the Toba-Qom communities, we selected the participants with data for at least 80% of the nights in the cycle. Sixty-nine participants met this condition: 20 in Ru-LL, 23 in Ru-NL, and 26 in Ur. Each participant’s data points were first averaged by night in the cycle (for those with data for over a whole cycle), and data for any missing nights were completed by linear interpolation. Data were then run through a moving-average filter with a 7-night window to eliminate low-frequency noise. Individuals’ data and whole-population data were fitted through a Non-linear Least Squares approach to the best possible sine curve with a 30-day period using the *nls* tool from the *stats* package (R Core Team 2018). Because recent literature has pointed to the possibility of semilunar month (~15 days) rhythms in mood and sleep^11–13^, alternative fits were performed considering the combination of a sine wave with a 30-day period along a 15-day period harmonic wave.

*Goodness-of-fit* evaluation was measured by the standard error of the regression (S), a proper measure for non-linear equation fits. When analysing group differences from data from individual fits we only included subjects within the best three quantiles of S values for their z-score fits for each variable, sleep duration and sleep onset (51 out of the 69 participants which data was fitted in each).

Rayleigh z-tests were used to analyse phase clustering and were performed and plotted with El Temps software v1.311 (University of Barcelona, Spain). Fiducial limits were used as a measure of confidence intervals of the mean phases, due to the non-normal nature of the circular distribution. Circular distributions were compared by Watson-Wheeler Tests for Homogeneity of phases using the *circular* package for R^24^.

Because we did not count with longitudinal recordings for each college student throughout the moon phases these data were analysed as follows. We first calculated the average and standard deviation of sleep duration on school nights (Sunday to Monday, excluding nights before a holiday) for each season. Then, we normalized each sleep event duration by subtracting the season average and dividing it by the season standard deviation. We then averaged these population data according to the phase of the moon cycle and smoothed it by running a 7-night moving-average. The data were then fitted to a sine wave with a 30-day period through the Non-linear Least Squares approach.

#### Statistical analysis

To estimate the associations between the moonlight availability phases and demographic variables on activity and sleep features we applied linear mixed-effects models (LMEM) using the *lme4* package^25^. Only participants who presented data for at least four nights within each moonlight phase were considered for these analyses. QQ-plots for every LMEM fit were visually inspected to check normality of residues and consider necessary transformation of variables; these are presented in the Extended Data. All the random effects (intercepts by subject) of the best-fitted models were normally distributed. The WASO variable required a square-root transformation to achieve linearity of residues. The syntax used for LMEMs is also presented in the Extended Data.

Cohen’s D effect size for each fixed effect of the best fitted models was calculated using the *lme.dscore* function of the *EMATools* R package^26^. This package calculates the Cohen’s D using the following equation:

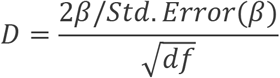

The degrees of freedom of the model (df) were calculated using the Satterthwaite approximation (via the *lmerTest* R library^27^). Confidence intervals for the fixed effect estimations were calculated using the *confint* function of the *stats* R library with alpha level set at 5% and using the Wald method. For the confidence intervals of the effect size we followed the *lme.dscore* methodology but using the beta estimates limits of the CI instead of the beta estimate itself.

## Author contributions

LC designed the study, collected Toba/Qom data, analyzed data, and wrote the manuscript. IS analyzed data and wrote the manuscript. GD collected Toba/Qom and UW students’ data. KMG organized and processed Toba/Qom recordings. EFD and CV provided resources for the study and wrote the manuscript. HOD designed the study, provided resources for the study, collected Toba/Qom and UW students’ data, analyzed data, and wrote the manuscript.

## Acknowledgments

We are thankful to Adán García, Mirtha Pérez, Karina Ortiz, Marcelo Rotundo, Ernesto Ruiz Guiñazú, Nastassya West for field assistance, and the Toba/Qom people for the tremendous support and unparalleled compliance. We thank Dr. William Scwartz for comments to the manuscript.

## Data availability

Toba/Qom data will be made openly available after publication. College students’ data will be made fully available after publication of a separate study from which these data was taken.

