## Extended Data for "Moonstruck sleep: Synchronization of Human Sleep with the Moon Cycle under Natural Conditions"

### Statistical model definitions

1) Comparing sleep duration and onset between communities

Syntax example of Linear-Mixed Effects Models used to analyse sleep variables between moonlight phases. The *lmer* function belongs to the *lme4 R* package[25]. "Fitting Linear Mixed-Effects Models Using lme4." Journal of Statistical Software, 67(1), 1–48)

*Model <- lmer(Variable ~ Group + Sex + Age + (1|ID),*
*data = dataframe)*

QQplots for the LMEMs presented in Extended Data Table 2

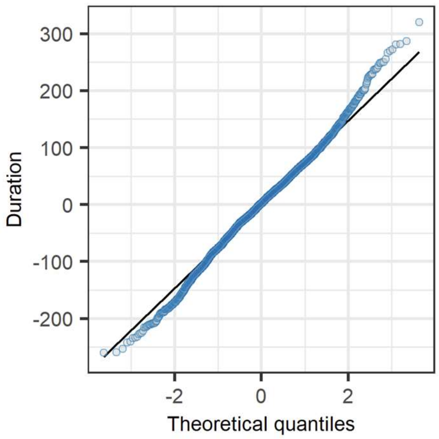


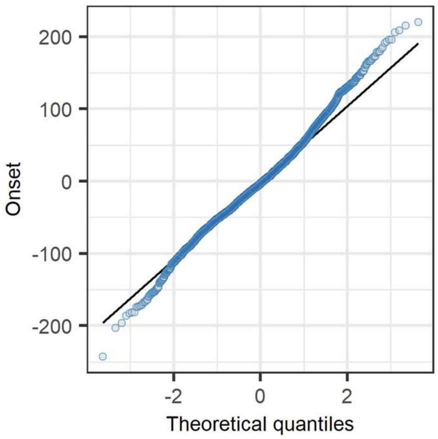

2) Comparing sleep variables between the Full Moonlight and No Moonlight phases

Syntax example:

*Model <- lmer(Variable ~ Moonlight_phase * Group + Sex + Age + (1|ID),*

*data = dataframe)*

<u>QQplots for the six LMEMs presented in full in Extended Data Table 4</u>

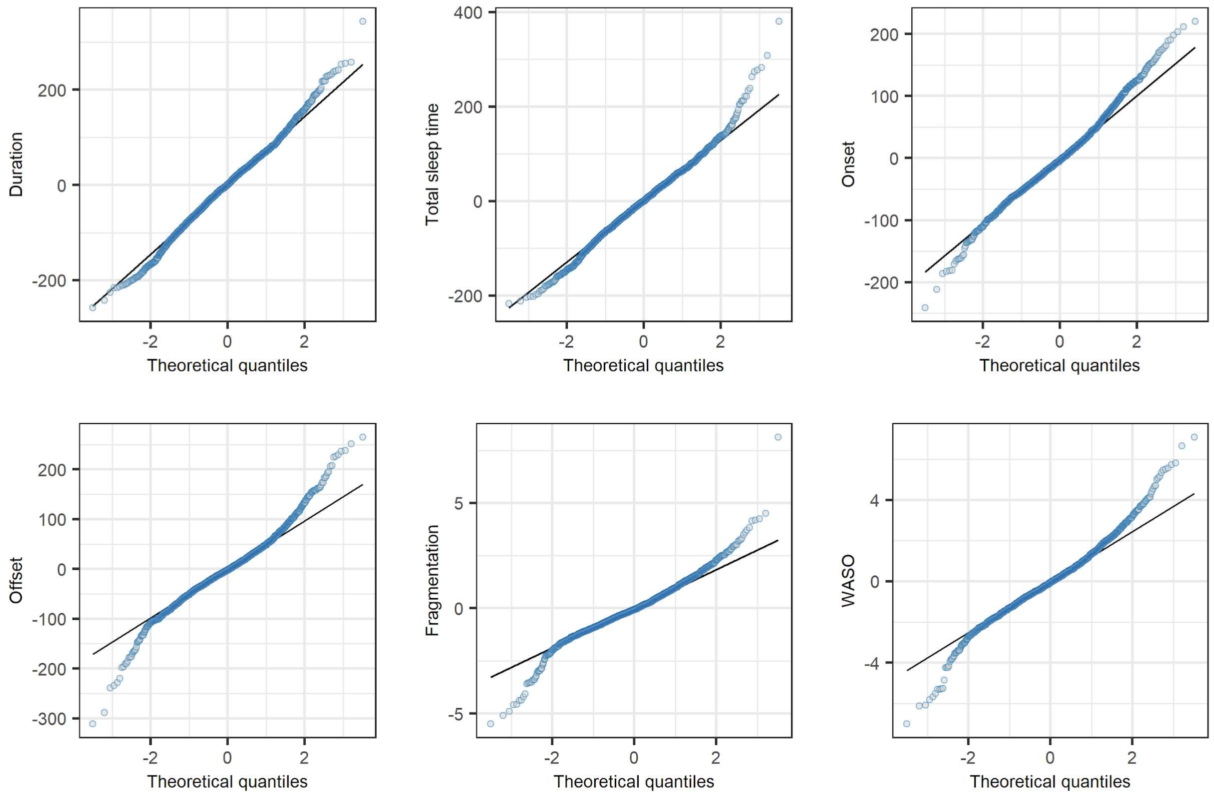

3) Comparing sleep offset between the between nights in which there is moonlight available for the last six hours before dawn (MLBD) and those without any moonlight in those six hours (No-MLBD)

Syntax example:

*Model <- lmer(Offset ~ Moonlight_phase * Group + Sex + Age + (1|ID),*

*data = dataframe)*

<u>QQplot for the LMEM presented in Extended Data Table 6</u>

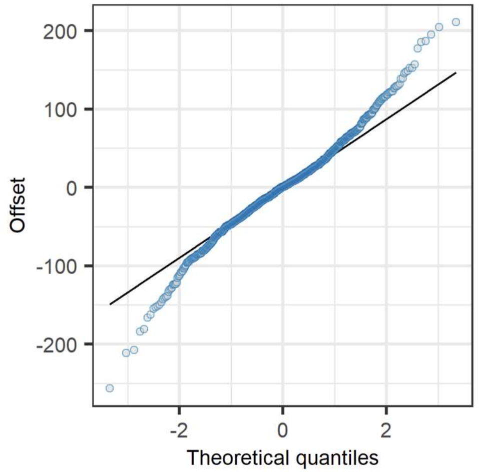

**Comparison of communities' phases from fitted data**

*1) Sleep duration through phases*

Mean in days to (negative values) or from (positive values) the full moon [fiducial limits]:

Rural, no light community: -1.5 [-3.4 / 0.4]

Rural, limited light: -3.1 [-5.8 / -0.4]

Urban: -6.6 [-8.5 / -4.7]

Watson-Wheeler Test for Homogeneity of phases between groups (from *circular* package in R):

W = 6.5348, df = 4, p = 0.1626

*2) Sleep onset peak phases*

Mean in days to (negative values) or from (positive values) the full moon [fiducial limits]:

Rural, no light community: -4.9 [-7.2 / -2.7]

Rural, limited light: -1.0 [-2.8 / 0.4]

Urban: -4.1 [-5.6 / -2.7]

Watson-Wheeler Test for Homogeneity of phases between groups:

W = 9.1582, df = 4, p = 0.0573

### Extended Data Tables

***Extended Data Table 1.*** *Composition of study communities*

| Group | n | Females (%) | Age mean (range) | Year(s) |
|---|---|---|---|---|
| Rural, no light | 25 | 15 (60%) | 23.2 (12 - 51) | 2016 |
| Rural, limited light | 33 | 17 (50%) | 22.7 (12 - 57) | 2016, 2018 |
| Urban | 40 | 23 (58%) | 25.8 (12 - 75) | 2017, 2018 |
| UW students | 563 | 353 (63%) | 21.4 (20.4 - 21.9) | 2015-2018 |

Note: n: number of participants

***Extended Data Table 2.*** *Fixed-effect terms included in the best-fitting model comparing sleep duration and onset of sleep between communities.*

| Magnitude | Fixed effect | Estimate | 95% CI | | Std. error | df | p-value | Estimate | 95% CI | |
|---|---|---|---|---|---|---|---|---|---|---|
| **Duration (min)** | (Intercept) | 516.13 | 498.55 | 533.74 | 9.14 | 81.42 | < 2e-16 | - | - | - |
| | Group - Ru-LL | 22.14 | 8.18 | 36.08 | 7.25 | 79.81 | 0.00308 | 0.683 | 0.253 | 1.114 |
| | Group - Ru-NL | 34.21 | 20.14 | 48.27 | 7.31 | 75.86 | 1.24E-05 | 1.074 | 0.632 | 1.515 |
| | Sex (Male) | 6.06 | -5.69 | 17.79 | 6.10 | 78.78 | 0.32425 | 0.224 | -0.210 | 0.657 |
| | Age | -1.95 | -2.52 | -1.39 | 0.29 | 88.20 | 2.67E-09 | -1.411 | -1.821 | -1.001 |
| **Onset (min from dusk)** | (Intercept) | 132.11 | 109.81 | 154.37 | 11.58 | 80.78 | <2e-16 | - | - | - |
| | Group - Ru-LL | -11.28 | -29.01 | 6.45 | 9.22 | 80.23 | 0.2246 | -0.273 | -0.703 | 0.156 |
| | Group - Ru-NL | -21.83 | -39.87 | -3.78 | 9.38 | 78.60 | 0.0226 | -0.525 | -0.959 | -0.091 |
| | Sex (Male) | 15.00 | 0.04 | 29.96 | 7.78 | 79.72 | 0.0573 | 0.432 | 0.001 | 0.863 |
| | Age | 0.72 | 0.01 | 1.43 | 0.37 | 83.27 | 0.0538 | 0.429 | 0.007 | 0.851 |

Note: These models were fit from data from all participants with records: Ru-NL, 25; Ru-LL, 29; Ur, 33

***Extended Data Table 3.*** *Descriptive statistics of sleep variables according to the No Moonlight (No-ML) and the Full Moonlight (F-ML) phases.*

| Group | n | Phase | Duration (min) | | | Sleep Time (min) | | | Fragmentation (wakes/hr of sleep) | | |
|---|---|---|---|---|---|---|---|---|---|---|---|
| | | | Mean | SD | 95%CI | Mean | SD | 95%CI | Mean | SD | 95%CI |
| Rural, no light | 25 | F-ML | 488 | 38 | 472 - 503 | 428 | 35 | 413 - 442 | 4.3 | 0.8 | 4 - 4.7 |
| | | No-ML | 513 | 35 | 499 - 527 | 449 | 33 | 436 - 463 | 4.3 | 0.8 | 3.9 - 4.6 |
| Rural, limited light | 25 | F-ML | 483 | 30 | 471 - 495 | 415 | 33 | 401 - 429 | 4.8 | 1 | 4.4 - 5.2 |
| | | No-ML | 504 | 38 | 488 - 520 | 437 | 37 | 421 - 452 | 4.5 | 0.9 | 4.2 - 4.9 |
| Urban | 32 | F-ML | 461 | 47 | 444 - 478 | 407 | 48 | 390 - 425 | 3.9 | 0.9 | 3.6 - 4.2 |
| | | No-ML | 472 | 46 | 455 - 489 | 417 | 45 | 401 - 433 | 3.9 | 0.8 | 3.6 - 4.2 |

| Group | n | Phase | Onset (min from dusk) | | | Offset (min from dawn) | | | WASO (min) | | |
|---|---|---|---|---|---|---|---|---|---|---|---|
| | | | Mean | SD | 95%CI | Mean | SD | 95%CI | Mean | SD | 95%CI |
| Rural, no light | 25 | F-ML | 153 | 48 | 133 - 172 | 85 | 46 | 66 - 104 | 60 | 12 | 55 - 65 |
| | | No-ML | 129 | 34 | 115 - 142 | 80 | 41 | 63 - 97 | 64 | 15 | 57 - 70 |
| Rural, limited light | 25 | F-ML | 157 | 36 | 142 - 172 | 116 | 49 | 95 - 136 | 68 | 16 | 61 - 75 |
| | | No-ML | 138 | 33 | 124 - 152 | 119 | 49 | 99 - 140 | 67 | 16 | 60 - 74 |
| Urban | 32 | F-ML | 164 | 46 | 147 - 181 | 112 | 57 | 91 - 132 | 54 | 14 | 49 - 59 |
| | | No-ML | 154 | 45 | 138 - 170 | 114 | 52 | 95 - 133 | 55 | 16 | 49 - 61 |

<u>Note:</u> The table contains data from the 82 participants who presented records for at least four nights in each moonlight phase. n: number of participants.

***Extended Data Table 4.*** *Fixed-effect terms included in the best-fitting model comparing sleep variables between the No Moonlight (No-ML) and the Full Moonlight (F-ML) phases.*

| | | **Fixed effects** | | | | | | **Effect size (Cohen d)** | | |
|---|---|---|---|---|---|---|---|---|---|---|
| **Magnitude** | **Fixed effect** | **Estimate** | **95% CI** | | **Std. error** | **df** | **p-value** | **Estimate** | **95% CI** | |
| **Duration (min)** | (Intercept) | 537.22 | 518.53 | 555.91 | 9.54 | 91.39 | < 2e-16 | - | - | - |
| | **ML-Phase (No-ML)** | **25.09** | **13.40** | **36.77** | **5.96** | **2110.26** | **2.67E-05** | **0.183** | **0.098** | **0.269** |
| | Group - Ru-LL | -7.56 | -25.08 | 9.95 | 8.94 | 112.21 | 0.3991 | -0.160 | -0.530 | 0.210 |
| | Group - Urban | -18.03 | -34.62 | -1.44 | 8.46 | 110.30 | 0.0353 | -0.406 | -0.779 | -0.033 |
| | Sex (Male) | 1.43 | -11.16 | 14.02 | 6.43 | 72.49 | 0.8243 | 0.052 | -0.408 | 0.513 |
| | Age | -2.17 | -2.77 | -1.57 | 0.31 | 82.12 | 4.74E-10 | -1.560 | -1.992 | -1.127 |
| | ML-Phase:Group - Ru-LL | -5.92 | -23.09 | 11.24 | 8.76 | 2131.05 | 0.4991 | -0.029 | -0.114 | 0.056 |
| | ML-Phase:Group Urban | -14.59 | -30.31 | 1.13 | 8.02 | 2119.01 | 0.0691 | -0.079 | -0.164 | 0.006 |
| **Total sleep time (min)** | (Intercept) | 473.29 | 453.81 | 492.77 | 9.94 | 90.10 | < 2e-16 | - | - | - |
| | **ML-Phase (No-ML)** | **22.03** | **11.54** | **32.52** | **5.35** | **2111.72** | **4.02E-05** | **0.179** | **0.094** | **0.264** |
| | Group - Ru-LL | -14.52 | -32.54 | 3.50 | 9.20 | 104.97 | 0.1174 | -0.308 | -0.691 | 0.074 |
| | Group - Urban | -12.38 | -29.47 | 4.71 | 8.72 | 103.41 | 0.1585 | -0.279 | -0.665 | 0.106 |
| | Sex (Male) | -4.79 | -18.12 | 8.54 | 6.80 | 75.82 | 0.4832 | -0.162 | -0.612 | 0.288 |
| | Age | -1.91 | -2.54 | -1.28 | 0.32 | 83.40 | 6.43E-08 | -1.300 | -1.729 | -0.871 |
| | ML-Phase:Group - Ru-LL | -1.71 | -17.13 | 13.72 | 7.87 | 2127.07 | 0.8285 | -0.009 | -0.094 | 0.076 |
| | ML-Phase:Group Urban | -12.48 | -26.60 | 1.65 | 7.21 | 2118.19 | 0.0836 | -0.075 | -0.160 | 0.010 |
| **Onset (min from dusk)** | (Intercept) | 123.22 | 99.96 | 146.49 | 11.87 | 80.76 | < 2e-16 | - | - | - |
| | **ML-Phase (No-ML)** | **-21.66** | **-30.32** | **-13.01** | **4.42** | **2107.55** | **1.01E-06** | **-0.214** | **-0.299** | **-0.128** |
| | Group - Ru-LL | 7.79 | -13.32 | 28.90 | 10.77 | 86.91 | 0.4716 | 0.155 | -0.265 | 0.576 |
| | Group - Urban | 10.68 | -9.38 | 30.73 | 10.23 | 86.12 | 0.2998 | 0.225 | -0.198 | 0.647 |
| | Sex (Male) | 16.28 | -0.02 | 32.58 | 8.32 | 74.34 | 0.0541 | 0.454 | -0.001 | 0.909 |
| | Age | 0.92 | 0.16 | 1.68 | 0.39 | 77.97 | 2.04E-02 | 0.536 | 0.092 | 0.980 |
| | ML-Phase:Group - Ru-LL | 0.08 | -12.67 | 12.82 | 6.50 | 2115.44 | 0.9904 | 0.001 | -0.085 | 0.086 |
| | ML-Phase:Group Urban | 12.28 | 0.62 | 23.94 | 5.95 | 2110.88 | 0.0391 | 0.090 | 0.005 | 0.175 |

*Extended Data Table 4 (continued).*

| | | Fixed effects | | | | | | Effect size (Cohen d) | | |
|---|---|---|---|---|---|---|---|---|---|---|
| **Magnitude** | **Fixed effect** | **Estimate** | **95% CI** | | **Std. error** | **df** | **p-value** | **Estimate** | **95% CI** | |
| **Offset (min from dusk)** | (Intercept) | 118.91 | 91.74 | 146.08 | 13.86 | 81.59 | 5.17E-13 | - | - | - |
| | **ML-Phase (No-ML)** | **-3.02** | **-11.85** | **5.82** | **4.51** | **2109.27** | **0.5039** | **-0.029** | **-0.114** | **0.056** |
| | Group - Ru-LL | 28.58 | 4.03 | 53.13 | 12.53 | 86.30 | 0.0250 | 0.491 | 0.069 | 0.913 |
| | Group - Urban | 36.71 | 13.37 | 60.05 | 11.91 | 85.67 | 0.0028 | 0.666 | 0.243 | 1.090 |
| | Sex (Male) | 16.36 | -2.78 | 35.49 | 9.76 | 76.58 | 0.0979 | 0.383 | -0.065 | 0.831 |
| | Age | -1.81 | -2.70 | -0.92 | 0.45 | 79.45 | 1.45E-04 | -0.896 | -1.336 | -0.456 |
| | ML-Phase:Group - Ru-LL | 5.21 | -7.81 | 18.22 | 6.64 | 2115.20 | 0.4332 | 0.034 | -0.051 | 0.119 |
| | ML-Phase:Group Urban | 4.48 | -7.43 | 16.39 | 6.08 | 2111.78 | 0.4611 | 0.032 | -0.053 | 0.117 |
| **Fragmentation (wakes/hr of sleep)** | (Intercept) | 4.45 | 3.95 | 4.95 | 0.25 | 81.30 | < 2e-16 | - | - | - |
| | **ML-Phase (No-ML)** | **-0.08** | **-0.24** | **0.09** | **0.08** | **2110.00** | **0.3501** | **-0.04** | **-0.126** | **0.045** |
| | Group - Ru-LL | 0.39 | -0.06 | 0.84 | 0.23 | 86.10 | 0.0903 | 0.369 | -0.053 | 0.791 |
| | Group - Urban | -0.37 | -0.79 | 0.06 | 0.22 | 85.40 | 0.0977 | -0.362 | -0.785 | 0.062 |
| | Sex (Male) | 0.49 | 0.14 | 0.84 | 0.18 | 76.30 | 0.0076 | 0.628 | 0.179 | 1.078 |
| | Age | -0.01 | -0.03 | 0.00 | 0.01 | 79.20 | 0.1506 | -0.326 | -0.766 | 0.114 |
| | ML-Phase:Group - Ru-LL | -0.17 | -0.40 | 0.07 | 0.12 | 2120.00 | 0.1760 | -0.059 | -0.144 | 0.026 |
| | ML-Phase:Group Urban | 0.03 | -0.19 | 0.25 | 0.11 | 2110.00 | 0.8150 | 0.010 | -0.075 | 0.095 |
| **WASO (min)** | (Intercept) | 7.84 | 7.28 | 8.40 | 0.29 | 82.60 | < 2e-16 | - | - | - |
| | **ML-Phase (No-ML)** | **0.20** | **-0.02** | **0.43** | **0.11** | **2110.00** | **0.0735** | **0.078** | **-0.007** | **0.163** |
| | Group - Ru-LL | 0.41 | -0.11 | 0.92 | 0.26 | 89.70 | 0.1251 | 0.327 | -0.087 | 0.742 |
| | Group - Urban | -0.43 | -0.92 | 0.05 | 0.25 | 88.80 | 0.0847 | -0.370 | -0.787 | 0.046 |
| | Sex (Male) | 0.38 | -0.01 | 0.77 | 0.20 | 75.20 | 0.0629 | 0.435 | -0.017 | 0.887 |
| | Age | -0.02 | -0.03 | 0.00 | 0.01 | 79.30 | 0.0824 | -0.395 | -0.835 | 0.045 |
| | ML-Phase:Group - Ru-LL | -0.26 | -0.59 | 0.06 | 0.17 | 2120.00 | 0.1148 | -0.069 | -0.154 | 0.017 |
| | ML-Phase:Group Urban | -0.17 | -0.47 | 0.13 | 0.15 | 2110.00 | 0.2765 | -0.047 | -0.133 | 0.038 |

Note: The models were fit with data from 82 participants who presented records for at least four nights in each moonlight phase: Ru-NL, 25; Ru-LL, 25; Ur, 32.

***Extended Data Table 5.*** *Descriptive statistics of sleep offset between nights in which there is moonlight available for the last six hours before dawn (MLBD) and those without any moonlight in those six hours (No-MLBD)*

| **Community** | **Participants** | **Phase** | **Mean (**min after dawn) | **SD** | **95%CI** |
|---|---|---|---|---|---|
| Rural, no light | 24 | No-MLBD | 77 | 29 | 65 - 89 |
| | | MLBD | 65 | 34 | 51 - 79 |
| Rural, limited light | 22 | No-MLBD | 109 | 40 | 91 - 127 |
| | | MLBD | 122 | 43 | 103 - 141 |
| Urban | 27 | No-MLBD | 108 | 53 | 87 - 129 |
| | | MLBD | 112 | 45 | 94 - 130 |

*Note:* The table contains data from 73 participants who presented records for at least four nights in each moonlight phase.

***Extended Data Table 6.*** *Fixed-effect terms included in the best-fitting model comparing times of sleep offset between nights in which there is moonlight available for the last six hours before dawn (MLBD) and those without any moonlight in those six hours (No-MLBD)*

| | **Fixed effects** | | | | | | **Effect size (Cohen d)** | | |
|---|---|---|---|---|---|---|---|---|---|
| **Fixed effect** | **Estimate** | **95% CI** | | **Std. error** | **df** | **p-value** | **Estimate** | **95% CI** | |
| (Intercept) | 78.83 | 47.72 | 109.93 | 15.87 | 77.15 | 3.99E-06 | - | - | - |
| **ML-Phase (No-MLBD)** | **11.35** | **-0.46** | **23.15** | **6.02** | **1140.53** | **0.0598** | **0.112** | **0.132** | **0.362** |
| Group - Ru-LL | 56.06 | 31.78 | 80.35 | 12.39 | 105.91 | 1.59E-05 | 0.879 | -0.393 | 0.156 |
| Group - Urban | 49.42 | 26.80 | 72.05 | 11.54 | 103.77 | 4.15E-05 | 0.841 | -0.589 | -0.025 |
| Sex (Male) | 14.96 | -2.64 | 32.56 | 8.98 | 67.45 | 0.1004 | 0.406 | -0.303 | 0.380 |
| Age | -0.81 | -1.88 | 0.25 | 0.54 | 67.41 | 0.1386 | -0.365 | -1.242 | -0.703 |
| ML-Phase:Group - Ru-LL | -25.18 | -42.03 | -8.34 | 8.60 | 1145.46 | 0.0035 | -0.173 | -0.159 | 0.077 |
| ML-Phase:Group Urban | -16.08 | -31.82 | -0.35 | 8.03 | 1144.77 | 0.0454 | -0.118 | -0.223 | 0.008 |

Note: The model was fit with data from 73 participants who presented records for at least four nights in each moonlight phase: Ru-NL, 24; Ru-LL, 22; Ur, 27.

### Extended Data Figures

#### Extended Data Figure 1

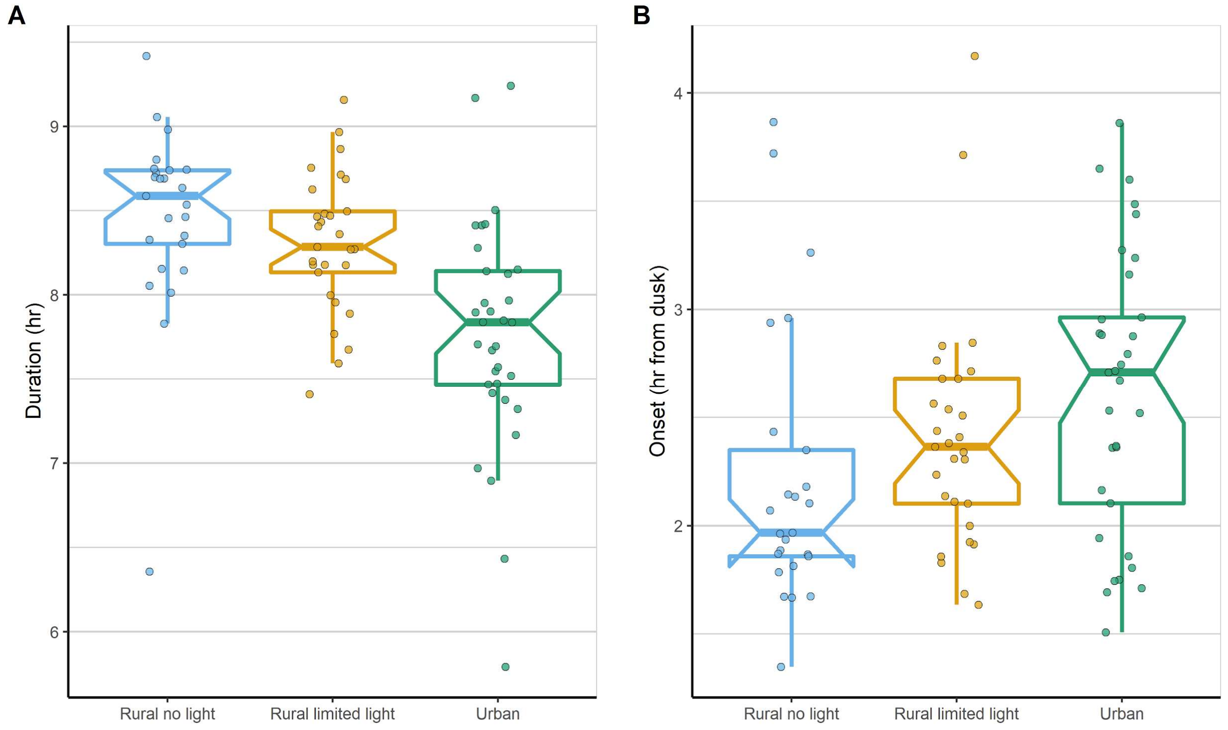


**Extended Data Figure 1. A)** Duration of sleep and **B)** time of sleep onset in the three Toba-Qom communities. Data summary is shown as boxplots with notches indicating the 95% CI of the median of the data. Dots represent individual participants' mean value for the variables across all the recording periods. LMEMs accounting for community, sex, and age as fixed effects, and subjects' identity as a random factor revealed associations between the level of access to electric light and both variables (see Extended Data Table 2). Participants per group: Ru-NL, 25; Ru-LL, 29; Ur, 33.

**Extended Data Figure 2**

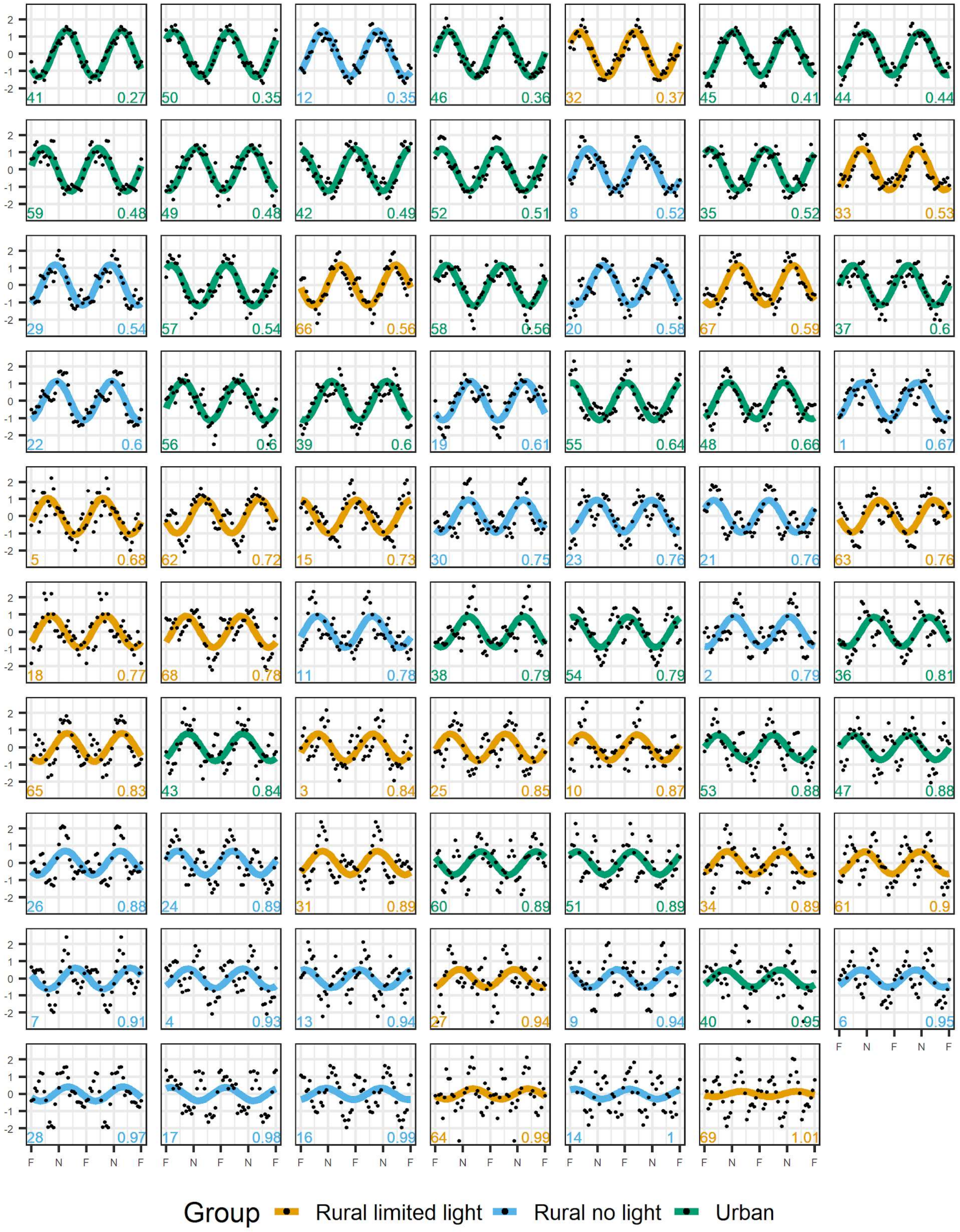


**Extended Data Figure 2.** Double-plots of sleep duration, expressed as *z*-scores, of individual participants in the study across the moon cycle. Dots indicate the duration of sleep on each night in the cycle, and coloured lines represent the best fit of a sine wave with a 30-days period through a non-linear least squares approach. The number on the bottom left of each plot identifies the participant and the one on the bottom right the standard error of the regression (S). The data correspond to participants with records for at least 80% of the moon cycle. Participants are ordered from best to worst fits as evaluated by S: minimum = 0.27, median = 0.76, maximum = 1.01. F = full moon, N = new moon.

### Extended Data Figure 3

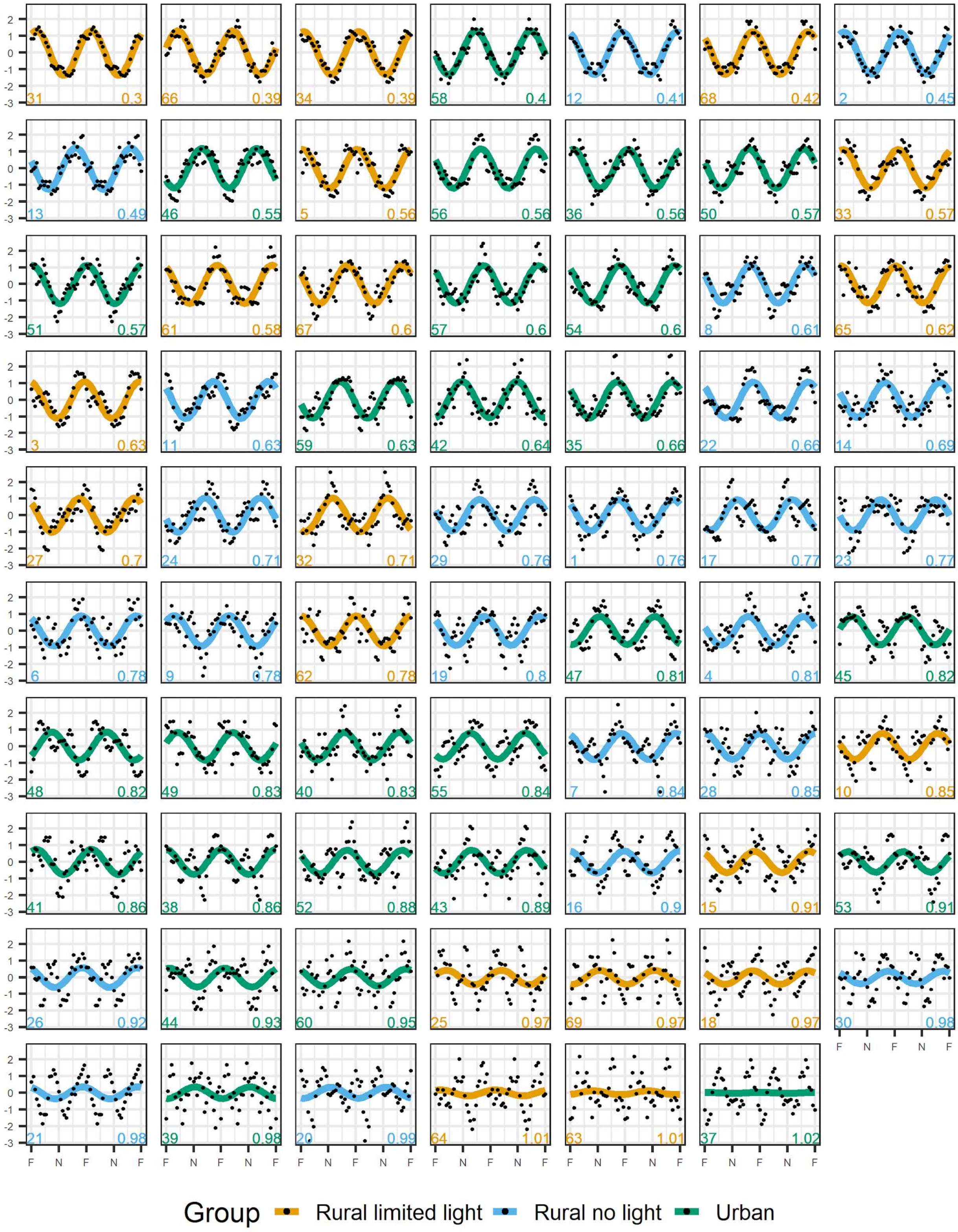


**Extended Data Figure 3.** Double-plots of sleep onset, expressed as *z*-scores, of individual participants in the study across the moon cycle. Dots indicate the onset of sleep on each night in the cycle, and coloured lines represent the best fit of a sine wave with a 30-days period through a non-linear least squares approach. The number on the bottom left of each plot identifies the participant and the one on the bottom right the standard error of the regression (S). The data correspond to participants with records for at least 80% of the moon cycle. Participants are ordered from best to worst fits as evaluated by S: minimum = 0.30, median = 0.78, maximum = 1.02. F = full moon, N = new moon.

**Extended Data Figure 4**

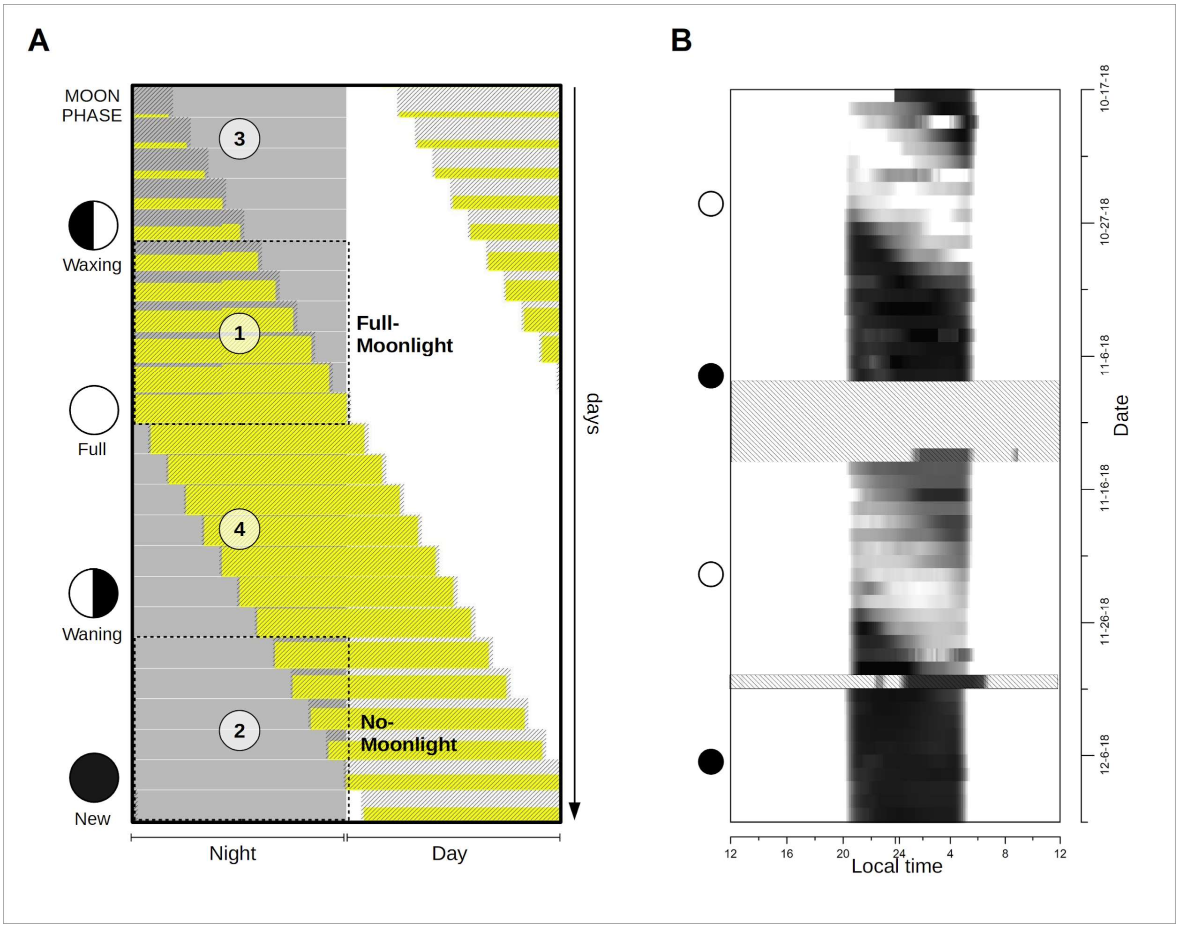


**Extended Data Figure 4**. **A)** Schematic representation of the progression of moonrise during a complete synodic moon cycle. Consecutive days are stacked vertically, with the shaded grey area representing the night hours. The textured areas represent the times when the moon is above the horizon, and the height of the yellow area represents the phase of the moon in terms of % of illuminated surface, from the new moon (no yellow) to the full moon (full yellow). Moonlight is available throughout the first six hours of the evening night in the interval indicated with number 1; we refer to it as the *Full Moonlight* phase (F-ML). During the interval indicated with number 2, no moonlight is available during said hours, and so we designate it as the *No Moonlight* phase (No-ML). The intervals indicated with numbers 3 and 4 are those in which the moon sets or rises during the first 6 hours of the evening, respectively. We refer to these generically as the *Waxing-Waning Moonlight* phases (W-ML). Note: for simplicity sake, the synodic lunar cycle was slightly shortened, and changes in the times of dusk and dawn were not simulated. **B)** Actual recording of sky illuminance from a location far from any electric light source near the Vaca Perdida settlement, from 10/17/2018 to 12/10/2018. The grayscale gradient was set so that the highest luminance levels equal to that of daylight. The brightest sky luminance recorded during the full moon was around 17.5 magnitudes per square arcsecond, while the darkest recordings obtained during new moons were around 22 mag/sq.arcsec, indicating a ~34-fold difference in night sky brightness between moon phases. Shaded boxes indicate missing or defective recording intervals. Data was recorded every 10 minutes with the Unihedron SQM-LU-DL device (Grimsby, ON, Canada).

### Extended Data Figure 5

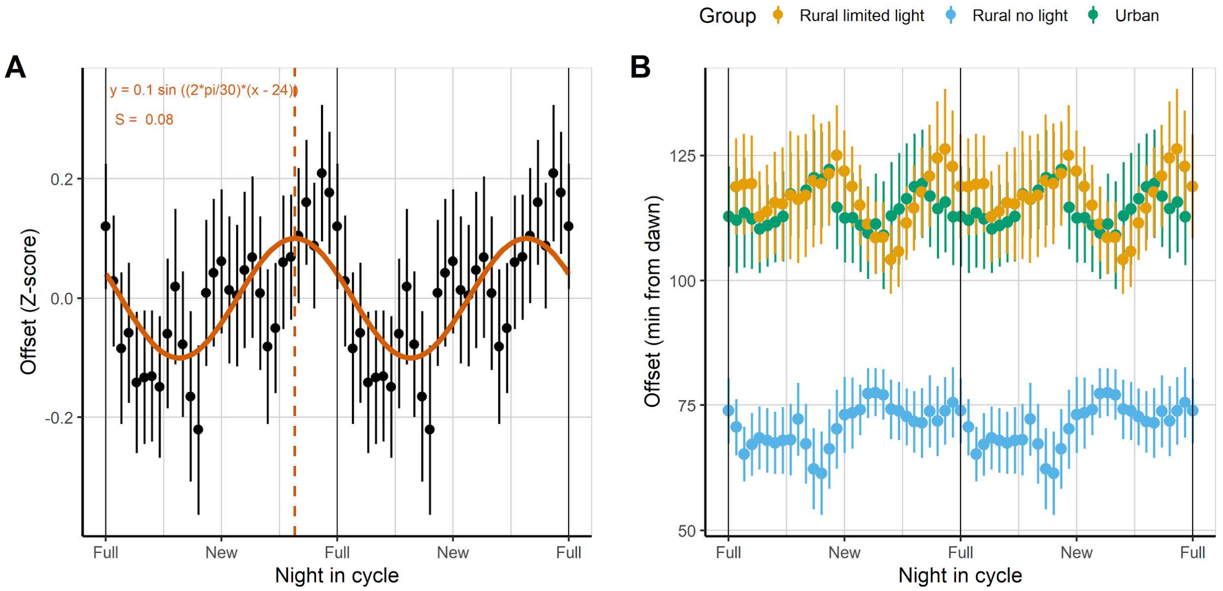


**Extended Data Figure 5.** Waking-up times across the moon cycle. **A)** Double-plot of the mean offset of sleep (expressed as z-scores) measured from the time of astronomical dusk across the moon cycle (N=69). The red solid line represents the best-fit to a sine curve with a 30-day period from a Non-Linear Least Squares fit, and the vertical dashed line indicates the acrophase of sleep offset (i.e. the latest waking-up time) according to the fit. The best-fit equation and the standard error of the regression (S) are indicated for the fit. **B)** Double-plot of the mean values of the offset of sleep in the three communities (Participants: Ru-LL, 20; Ru-NL, 23; Ur, 26). Vertical error bars on data points represent standard errors of the mean. Individual data series were filtered through moving-average with a window of seven days before summarizing the data. All data in these figures correspond to participants with records for at least 80% of the moon cycle.

**Extended Data Figure 6.**

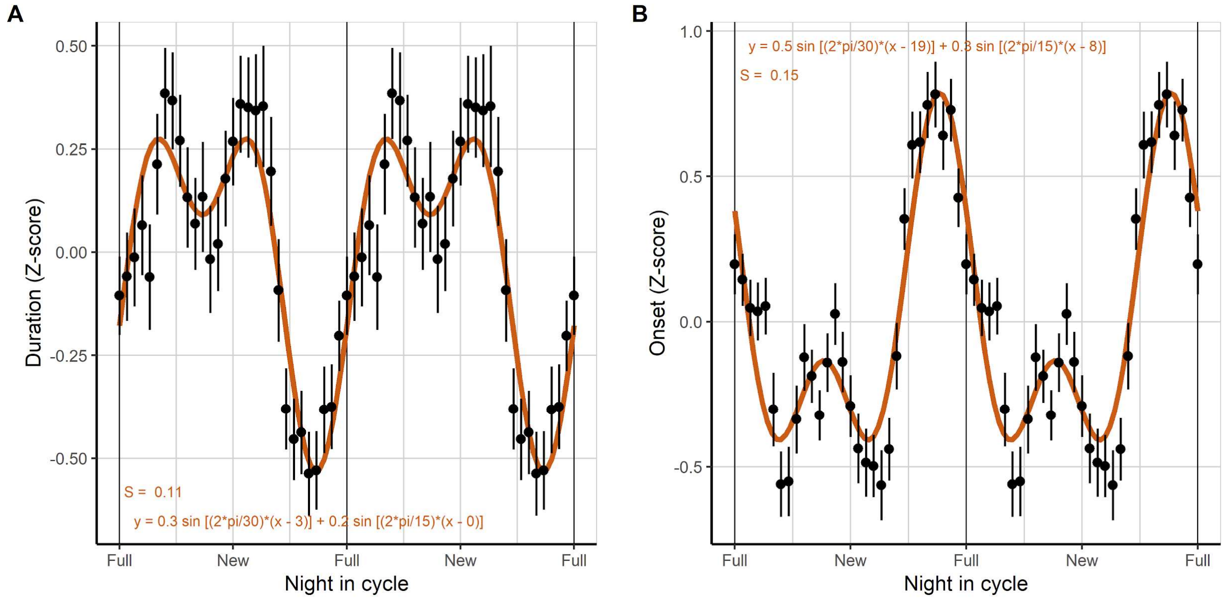


**Extended Data Figure 6.** Double-plots of **A)** the average duration and **B)** onset of sleep (measured from the time of astronomical dusk), expressed as z-scores (N=69). Red solid lines represent the best-fits to equations involving a sine curve with a 30-day period and a second sine curve with a 15-day period, calculated using a Non-Linear Least Squares approach. Individual data series were filtered through moving-average with a window of seven days before summarizing the data. All data in these figures correspond to participants with records for at least 80% of the moon cycle. The equation and standard error of the regression (S) are indicated for each fit.

**Extended Data Figure 7**

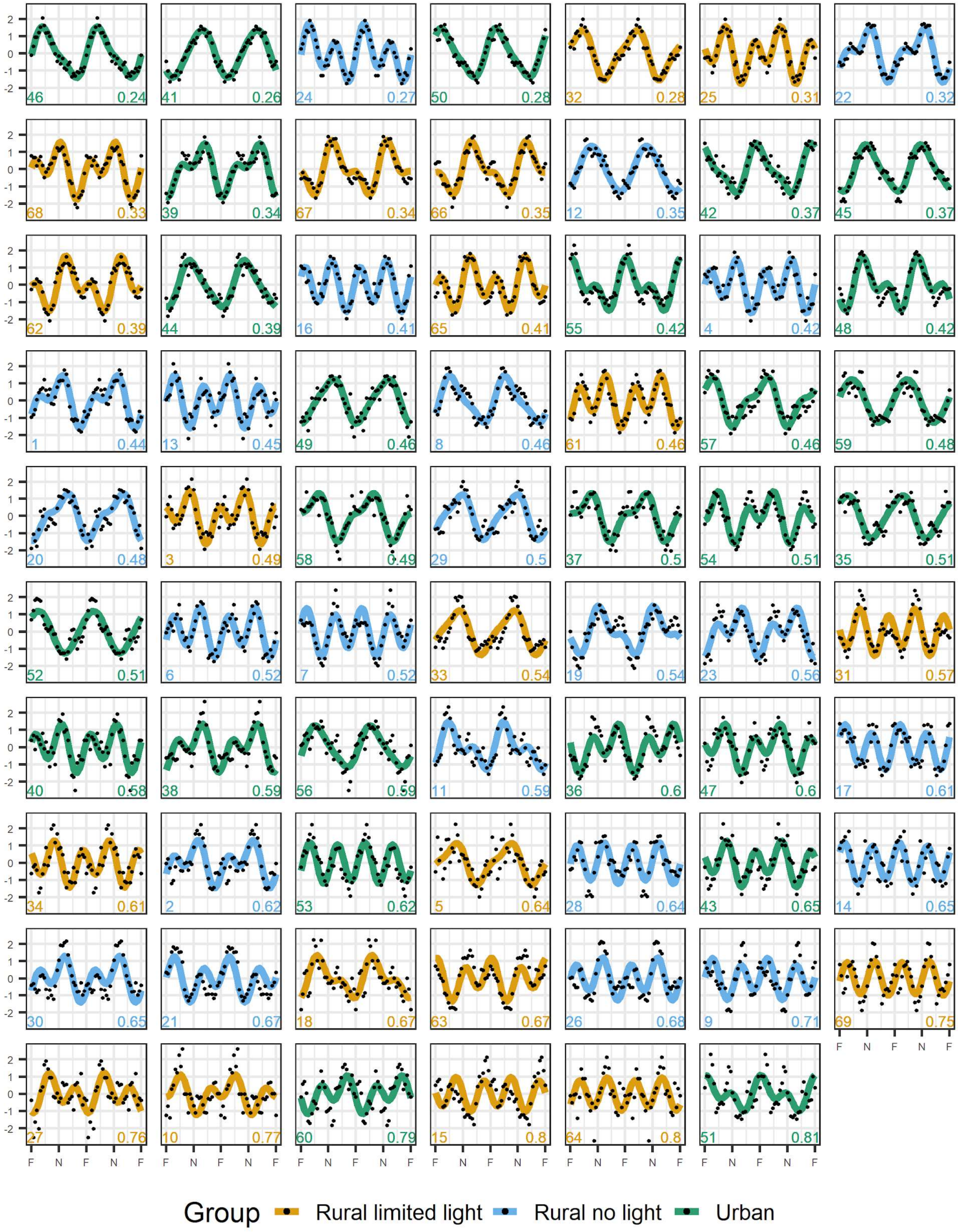


**Extended Data Figure 7.** Double-plots of sleep duration, expressed as *z*-scores, of individual participants in the study across the moon cycle. Dots indicate the duration of sleep on each night in the cycle, and coloured lines represent the best fit to an equation involving a sine curve with a 30-day period and a second sine curve with a 15-day period through a non-linear least squares approach. The number on the bottom left of each plot identifies the participant and the one on the bottom right the standard error of the regression (S). The data correspond to participants with records for at least 80% of the moon cycle. Participants are ordered from best to worst fits as evaluated by S: minimum = 0.24, median = 0.51, maximum = 0.81. F = full moon, N = new moon.

**Extended Data Figure 8**

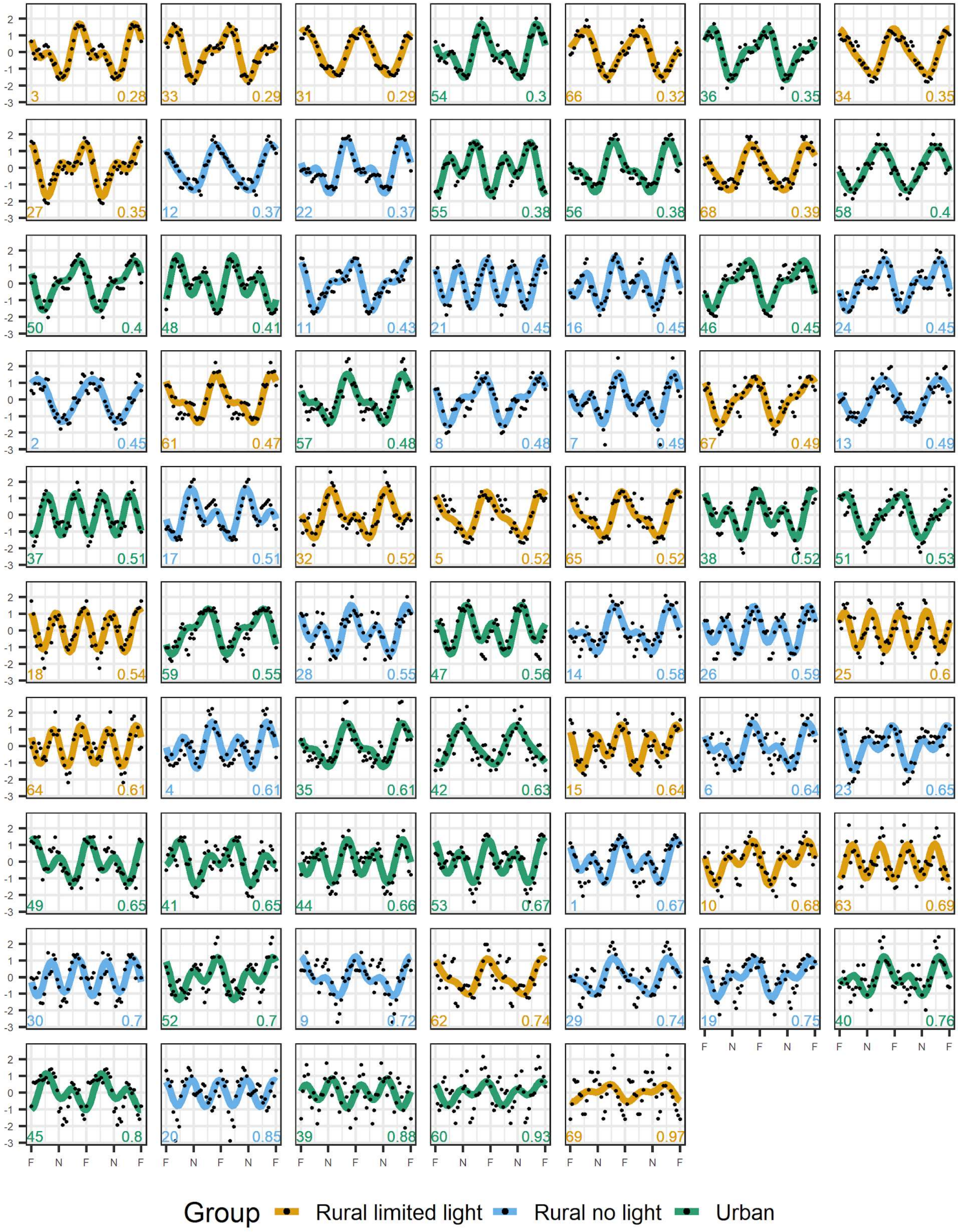


**Extended Data Figure 8.** Double-plots of sleep onset, expressed as *z*-scores, of individual participants in the study across the moon cycle. Dots indicate the duration of sleep on each night in the cycle, and coloured lines represent the best fit to an equation involving a sine curve with a 30-day period and a second sine curve with a 15-day period through a non-linear least squares approach. The number on the bottom left of each plot identifies the participant and the one on the bottom right the standard error of the regression (S). The data correspond to participants with records for at least 80% of the moon cycle. Participants are ordered from best to worst fits as evaluated by S: minimum = 0.28, median = 0.52, maximum = 0.97. F = full moon, N = new moon.
